# Antigen receptor locus dynamics is orchestrated near the sol-gel phase transition to enforce stepwise VDJ gene rearrangement

**DOI:** 10.1101/441444

**Authors:** Nimish Khanna, Yaojun Zhang, Joseph S. Lucas, Olga K. Dudko, Cornelis Murre

**Affiliations:** Division of Biological Sciences, 0377, Department of Molecular Biology, University of California, San Diego, La Jolla, CA 92093; Princeton Center for Theoretical Science, Princeton University, Princeton, NJ 08544; Department of Physics, University of California, San Diego, La Jolla, CA 92093

## Abstract

Diverse antibody repertoires are generated through remote genomic interactions involving immunoglobulin variable (V_H_), diversity (D_H_) and joining (J_H_) gene segments. How such interactions are orchestrated remains unknown. We developed a novel strategy to track V_H_-D_H_J_H_ motion and interactions in live B-lymphocytes. We found that V_H_ and D_H_J_H_ segments were trapped in configurations that only allowed constrained local motion, such that spatially proximal V_H_ and D_H_J_H_ segments remained in proximity, whereas spatially remote segments explored their immediate neighborhood while remaining remote. Comparison of experimental and simulated data revealed that such a highly constrained motion was imposed by a network of cross-linked chromatin chains characteristic of a gel phase, yet it was poised near the sol phase, a solution of independent chromatin chains. We propose that epigenetically induced gel droplets and the proximity to the sol-gel phase transition constitute the mechanism that orchestrates ordered VDJ rearrangement.

---

It is now well established that the mammalian genome, although dynamic, is highly structured. Chromosomes occupy individual territories and interact sparsely. Within chromosomes, chromatin loop domains are anchored by CTCF proteins and serve as the fundamental units of chromosome structure^1,2^. The folding of the loop domains is orchestrated by members of the cohesin complex, whereby cohesin extrudes loops of chromatin in a progressive fashion until a pair of convergent CTCF sites is reached^3^. The chromatin fiber is also subject to large-scale epigenetic modifications that include DNA methylation, chromatin remodelers, histone modifications and non-coding transcription. Recent studies have proposed that weak multivalent interactions result in phase-separated regions that compartmentalize gene expression^4^.

The immunoglobulin heavy chain (Igh) locus is assembled through long-range genomic interactions involving variable (V_H_), diversity (D_H_), joining (J_H_), and constant (C_H_) coding elements^5^. Two endonucleases, RAG1 and RAG2, initiate the recombination reaction by generating double-strand DNA breaks in the recombination signal sequences that flank the V_H_, D_H_ and J_H_ segments^6,7^. The variable regions are segregated into two distinct clusters, the distal and proximal V_H_ regions, that collectively span approximately 2.7 Mb. The D_H_ and J_H_ elements are separated from each other by 50 kb and are located upstream of the intronic enhancer Eμ. Located downstream of D_H_ and J_H_ are the constant regions as well as a cluster of CTCF sites named the Igh 3’ regulatory region, also known as the Igh super-anchor^5,8^. The V_H_ regions are associated with CTCF binding sites, which are positioned in a convergent orientation towards the super-anchor^8,9^. Two CTCF sites separate the V_H_ from the D_H_J_H_ elements to suppress predominant rearrangement of the proximal V_H_ region cluster and to orchestrate ordered Igh locus rearrangement^9,10,11^. Prior to V_H_-D_H_J_H_ rearrangement, the D_H_J_H_ region folds into a singular loop domain that insulates RAG activity across loop domains^12^.

The Igh locus is also characterized by a large spectrum of epigenetic modifications distributed across the V_H_ and D_H_J_H_ regions^13^. A subset of these modifications is deposited in a lineage- and developmental stage-specific fashion and correlate with ordered Igh V_H_-D_H_J_H_ rearrangement^14^. Prior to D_H_J_H_ rearrangement, non-coding transcription is initiated, and histone marks such as H3K27Ac, H3Ac and H3K4me1 are deposited within the D_H_J_H_ loop domain^15^. The deposition of these marks is controlled, at least in part, by an intronic enhancer located immediately downstream of the D_H_J_H_ region^13^. V_H_-D_H_J_H_ rearrangement is also associated with the deposition of epigenetic marks across the V_H_ regions that include H3Ac, H3K27me3, H3K4me1 and H3K27me3^16^. Finally, the chromatin remodeler BRG1 binds to multiple sites to assemble crosslinks across the Igh locus chromatin fiber, leading to Igh locus contraction and distal Ig V_H_-D_H_J_H_ rearrangement^17^.

Recent studies using a single-color labeling strategy demonstrated that V_H_ and D_H_J_H_ elements undergo anomalous diffusion in a viscoelastic environment whereby they bounce back and forth in a spring-like fashion ^18,19^. However, since only a single element was marked, these studies did not permit the visualization and description of V_H_-D_H_J_H_ genomic interactions and V_H_-D_H_J_H_ motion in live B cells. Here, we designed and generated tandem arrays of wild-type TET-operator and mutant TET-operator binding sites to separately mark both the V_H_ and the D_H_J_H_ elements. We found that the Igh locus adopted a wide spectrum of configurations that were dynamic enough to allow local motion yet stable enough to provide a strong long-term confinement. To determine the mechanistic origin of this confined motion, we performed Molecular Dynamics simulations of a hierarchy of polymer models. We found that that the Igh loop domains are organized as gel droplets poised near a sol-gel phase transition. We propose that the proximity of the Igh locus to the sol-gel phase boundary provides a tradeoff between stability and responsiveness: the ordered nature of the crosslinked network, formed by the locus and characteristic of a gel, enables ordered Igh locus rearrangement by facilitating and insulating V_H_-D_H_J_H_ and D_H_-J_H_ interactions, while the disordered nature of a solution of independent chains, characteristic of a sol, permits randomness in V_H_-D_H_J_H_ encounters and rapid Igh locus re-assembly.

## Results

### An approach to monitoring genomic interactions in live cells

In previous studies we tracked either V_H_ or D_H_J_H_ motion in live pro-B cells using tandem arrays of TET-operator binding sites^19^. We next aimed to monitor genomic encounters directly in live cells using a dual labeling approach (Fig. 1a). Attempts to insert tandem arrays of LAC operators in conjunction with TET-operator binding sites failed because tandem arrays of LAC operator binding sites were unstable in expanding pro-B cell cultures (data not shown). We therefore turned to a new strategy to simultaneously track the motion of paired genomic elements. Specifically, we used wild-type and mutant TET-repressor binding sites associated with unique DNA binding specificities^20,21^. From these, we selected a mutant TET repressor (4C5G) carrying two amino acid substitutions (E37A and P39K) in the TET-DNA binding domain since it showed nonoverlapping DNA binding activity (Fig. 1b). Tandem arrays containing 320 copies of the mutant TET 4C5G binding site (CCGTCAGTGACGG) were constructed essentially as described previously (Fig. 1c)^22^.

**Fig. 1.**
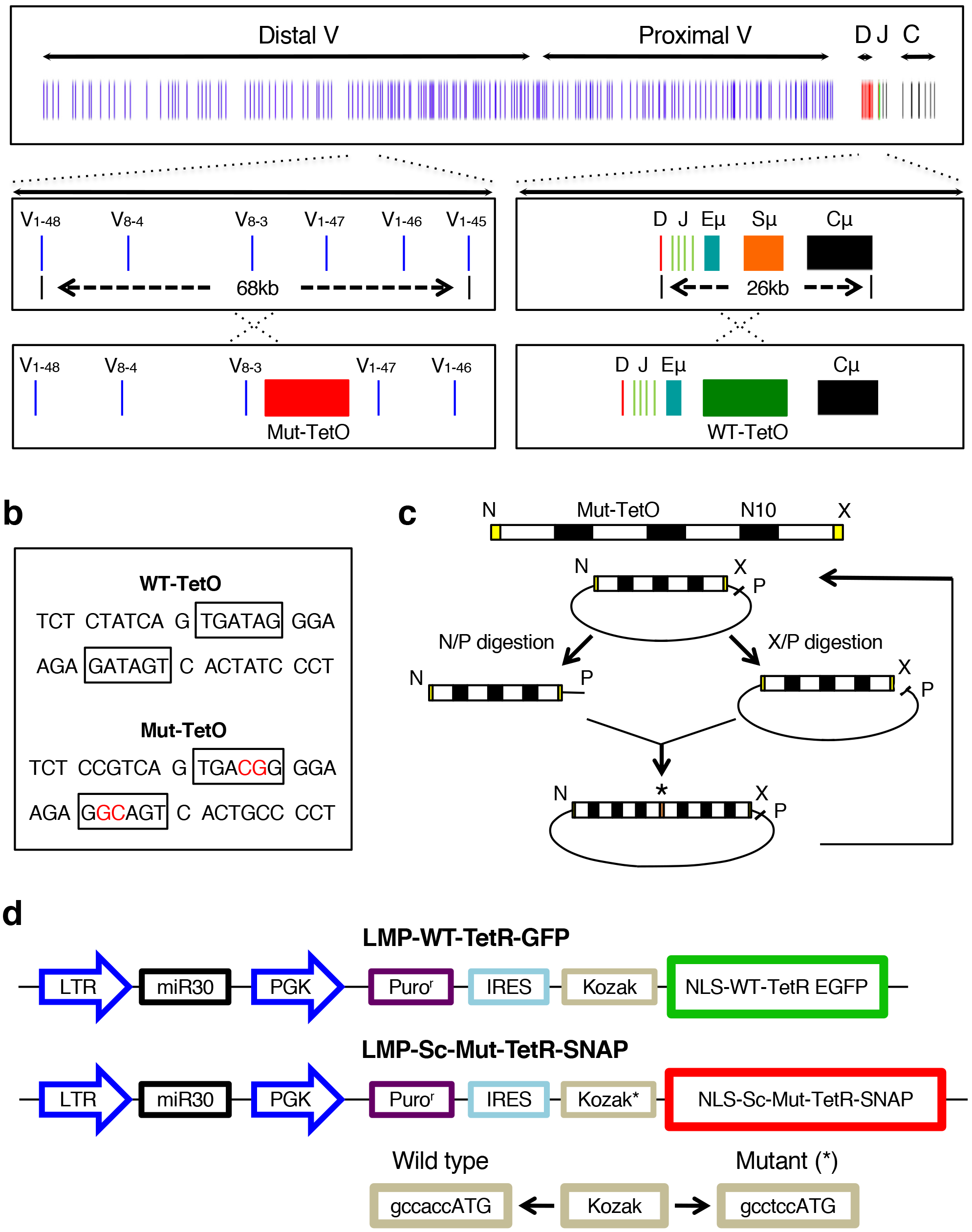
A Strategy to track V_H_-D_H_J_H_ motion in live B cells. **a**, Schematic of the Igh locus showing the V_H_, D_H_, J_H_, C_H_ segments, intronic enhancer (Eμ) and switch region repeats (Sμ). **b**, DNA sequences of wild-type and mutant TET operator binding sites. Nucleotides for which the mutant binding sites differ from that of WT-TET binding sites are indicated in red. **c**, Experimental strategy to design tandem arrays of mutant TET-operator binding sites. * indicates the junction of ligated DNA segments, each containing three MUT-TET binding sites. **d,** Construction of retroviral vectors to optimize the signal to noise ratio in pro-B cells transduced with virus expressing wild-type TET-EGFP and MUT-TET SNAP Tag.

In principle, two distinct repressor modules should permit the simultaneous tracking of two genomic regions. However, TET repressor binds as dimers and heterodimers of wild-type TET, and mutant TET protomers, which would interfere with the ability to bind tandem arrays of either WT-TET or MUT-TET binding sites specifically and selectively. To overcome this obstacle, we converted the MUT-TET repressor dimer into a single chain using a 29 amino acid linker. The single chain (sc) TET-repressor binds to its cognate binding site as a monomer^21^. To generate two differentially labeled fusion proteins, wild-type TET-repressor was fused to GFP (Fig. 1d). The mutant (sc) TET (4C5G) repressor was tagged with a polypeptide (SNAP-TAG) that has the ability to bind cell-permeable fluorescent dyes (Fig. 1d). To boost expression of the wild-type and mutant TET-repressors, both proteins were codon-optimized. Finally, to optimize the signal-to-background fluorescence ratio, the TET-SNAP-tag repressor was placed under control of a mutant Kozak sequence (GCCTCCATG) to damp translation and reduce background fluorescence (Fig. 1d).

### Tracking V_H_-D_H_J_H_ motion and V_H_-D_H_J_H_ encounters in live B-lymphocytes

In previous studies we have generated mice that carried tandem arrays of WT-TET operator binding sites in either the V_H_ or D_H_J_H_ regions^19^. From mice harboring a tandem array of WT-TET binding sites adjacent to the D_H_J_H_ region we derived immortalized RAG-deficient pro-B cell lines. Briefly, RAG1-deficient WT-TET D_H_J_H_ pro-B cells were isolated, grown in the presence of IL7 and SCF, transduced with BCR-ABL virus and cultured in methylcellulose-containing media. After seven days, colonies appeared that were expanded in the presence of S17 feeder cells, IL7 and SCF. This approach yielded BCR-ABL-transformed pro-B cell lines carrying a tandem array of TET-operator binding sites that marked the D_H_J_H_ region (Fig. 1a). Next, we inserted a tandem array of mutant TET repressor binding sites within the V_H_ region cluster. A template containing 320 copies of the mutant TET 4C5G binding site (CCGTCAGTGACGG) was constructed (Fig. 1c). The tandem array was next inserted into the V_H_ 8-3 region of D_H_J_H_ WT-TET pro-B cells using CAS9-CRISPR engineering. Stable clones were isolated using blasticidin as a selectable marker, expanded and examined for proper insertion into the genome using PCR and the appropriate. Two clones were isolated that contained the tandem array of MUT-TET binding sites inserted into the V_H_ region, expanded and transduced with virus expressing WT-TET GFP and MUT-TET SNAP-TAG (Fig. 1d). Two days post transduction, the cells were immobilized on poly-lysine coated dishes, incubated with the cell-permeable dye TMR-STAR to label MUT-TET SNAP-TAG bound sites and imaged using the OMX microscope platform. Cells transduced with virus expressing WT-GFP revealed two fluorescent foci consistent with a tandem array of WT-TET binding sites inserted near the D_H_J_H_ region on both alleles (Fig. 2a; left panels). Cells transduced with virus expressing MUT-TET SNAP tag revealed a singular fluorescent center consistent with the insertion of a MUT-TET tandem array in the distal V_H_ region on a single allele (Fig. 2a). Transduced V_H_ MUT-TET; D_H_J_H_ WT-TET pro-B cells were imaged every 2 seconds for 400 seconds in short term imaging or every 40 seconds for 4800 seconds (Supplementary Movies 1 and 2).

**Fig. 2.**
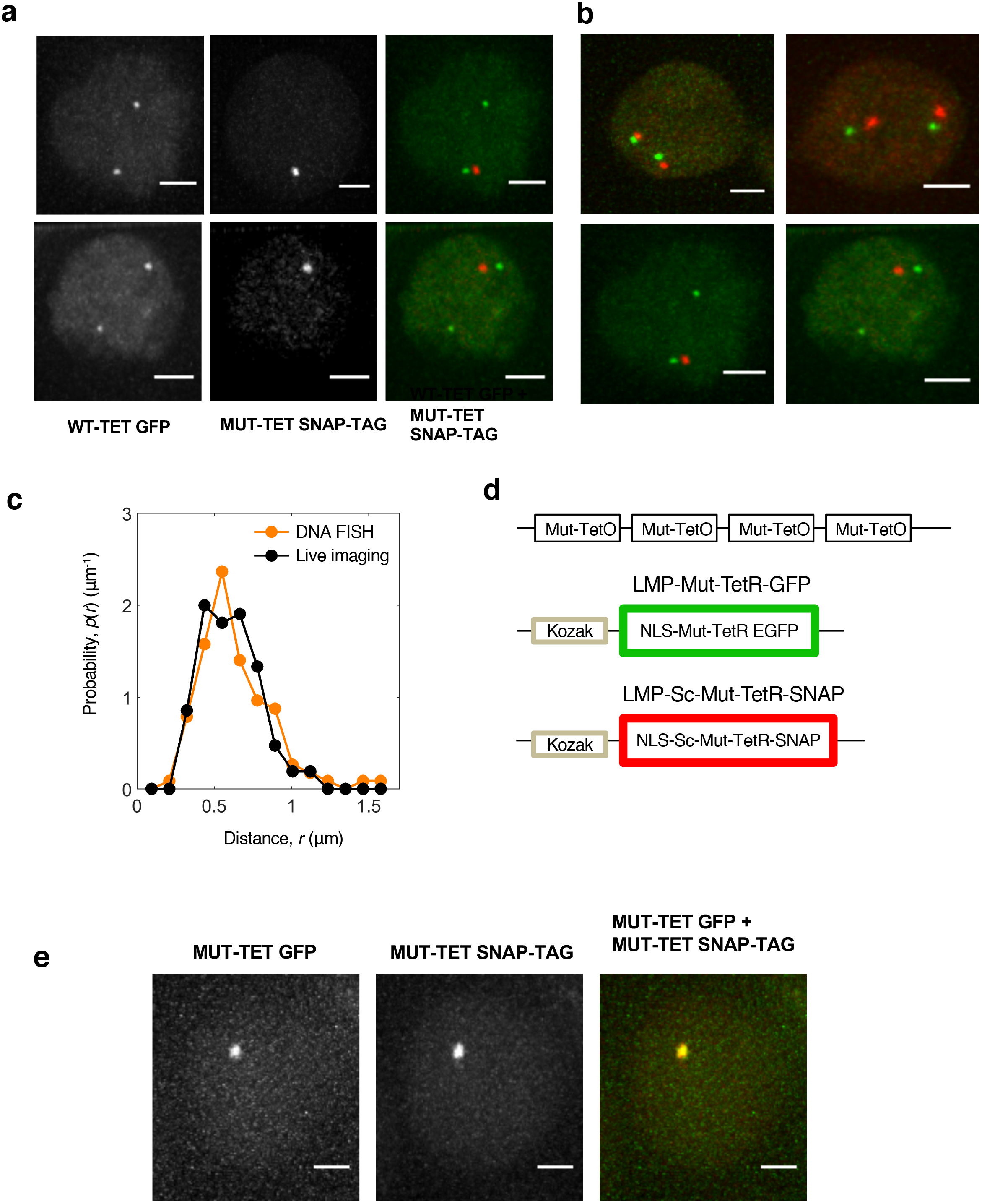
Tracking V_H_-D_H_J_H_ Motion in Pro-B Cells. **a**, Live imaging of pro-B cells harboring D_H_J_H_ WT-TET and V_H_ MUT-TET alleles. Panels indicate pro-B cells carrying D_H_J_H_ WT-TET and V_H_ MUT-TET alleles transduced with WT-TET GFP and MUT-TET SNAP-TAG. Top and middle panels show fluorescent foci in cells transduced with virus expressing WT-GFP and MUT-SNAP-TAG. Right-hand images show both green and red fluorescent foci representing labeled D_H_J_H_ and V_H_ regions. **b**, Pro-B cell lines harboring tandem arrays of WT-TET GFP and MUT-TET SNAP-TAG were analyzed by 3D-FISH using fluorescently labeled V_H_ and D_H_J_H_ probes (top panel) or obtained from fluorescently marked TET probes during live imaging (lower panel) for the same cell population. **c**, Distributions of spatial separations of the V_H_ and D_H_J_H_ elements obtained from live pro-B cell imaging (black dots) compared to those obtained from 3D-FISH measurements (orange dots). **d**, Experimental strategy to determine localization error. Localization error was determined by measuring the spatial distances separating red and green fluorescent signals.

To validate the experimental strategy using wild-type and mutant operator sites as a means to track long-range genomic interactions, we compared the distribution of the spatial distances separating the V_H_ and D_H_J_H_ elements, obtained by tracking V_H_-D_H_J_H_ motion in cells carrying WT-TET and MUT-TET binding sites, to those obtained by 3D-FISH measurements (Fig. 2b). We found that both methods yielded similar distributions, consistent with the insertion of tandem arrays of WT-TET and MUT-TET binding sites into the V_H_ and D_H_J_H_ regions, respectively (Fig. 2c). To determine the localization error associated with marking paired genomic regions with fluorescently labeled wild-type and mutant TET proteins, pro-B cells harboring tandem arrays of MUT-TET binding sites were transduced with virus expressing MUT-TET GFP (green fluorescence) and virus expressing MUT-TET SNAP-TAG (red fluorescence) (Fig. 2d). Cells that showed overlapping green and red fluorescent foci were imaged every 2 seconds for 400 seconds. The positions of the two fluorescent foci as a function of time were recorded to determine localization error. We found mean-squared displacement due to the measurement error to be approximately 0.02 μm^2^, or an average localization error of 22 nm for the *x*- and *y*-coordinates and 95 nm for the *z*-coordinate for each marker (Supplementary Information). To extract the true V_H_-D_H_J_H_ genomic motion from the observed apparent motion, the rotational and translational nuclear motion as well as localization error were eliminated using two independent procedures (Supplementary Information), neither one involving any adjustable parameters and both found to yield very similar outcomes (Supplementary Fig. 1). Taken together, these results showed that, by using tandem arrays of wild-type and mutant TET operator binding sites to mark coding or regulatory DNA elements, paired remote genomic interactions can be tracked with relatively high accuracy.

### V_H_-D_H_J_H_ trajectories reveal highly constrained, local motion

The individual trajectories of the spatial distances separating the labeled V_H_ and D_H_J_H_ regions revealed that V_H_-D_H_J_H_ motion was extremely confined: the V_H_ and D_H_J_H_ segments that were initially in close spatial proximity remained in proximity, whereas the segments that were initially spatially remote remained remote (Fig. 3a and Supplementary Fig. 2). The corresponding probability distribution of V_H_-D_H_J_H_ distances generated from these trajectories covered a broad range of distances (0.2-1.2 μm) suggesting that, across the population of cells, the locus adopts a wide spectrum of chromatin configurations (Fig. 3a; right panel). The distribution of V_H_-D_H_J_H_ distances exhibited bimodality that persisted for at least 400 seconds, indicating at least two dominant types of Igh locus configurations. Color-coding the trajectories of V_H_-D_H_J_H_ spatial distances according to their mean values revealed an intriguing de-mixing effect: the distances within the individual pairs fluctuated around respective mean values that remained nearly constant, together resulting in a “rainbow” pattern that persisted for at least an hour (Fig. 3a, left panel, and Supplementary Fig. 2). These data indicate that the Igh locus adopts a wide spectrum of configurations that are sufficiently dynamic to allow local motion, yet stable enough to provide a long-term (>60 minutes) confinement that either co-localizes V_H_ and D_H_J_H_ gene segments or isolates them from each other.

**Fig. 3.**
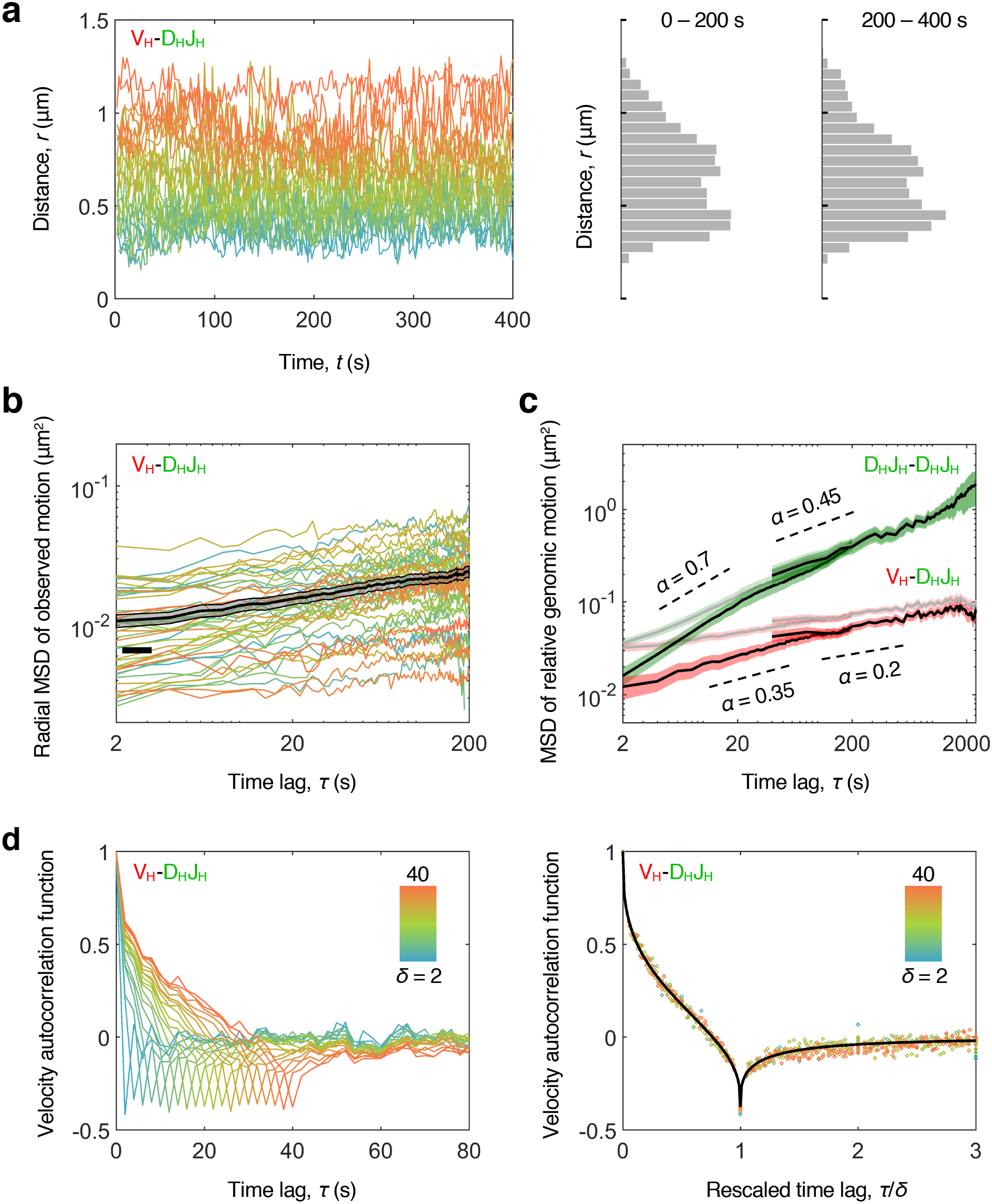
V_H_-D_H_J_H_ motion in B cell progenitors is extremely confined. **a,** The spatial distances within V_H_-D_H_J_H_ pairs fluctuates around respective average values that remain nearly constant over time. Color-coding the distance trajectories according to their mean values reveals a de-mixing effect. Right panel: the corresponding probability distribution of V_H_-D_H_J_H_ distances. **b**, The time-averaged radial MSD (colored lines) and the time-and-ensemble-averaged radial MSD (black line) for V_H_-D_H_J_H_ motion. **c,** Time-and-ensemble-averaged MSD of relative genomic motion for inter-chromosomal D_H_J_H_-D_H_J_H_ pair (green) and intra-chromosomal V_H_-D_H_J_H_ pair (red) before (pale lines) and after (bright lines) measurement-error correction. **d,** VAFs for the V_H_-D_H_J_H_ motion exhibit anti-correlations indicative of a push-back from the environment; the value at the dip approaches the theoretical limit (−0.5) of extreme confinement. The collapse of the VAF curves upon rescaling of the time axis (right panel) indicates self-similarity of motion.

The properties of V_H_-D_H_J_H_ motion averaged across the trajectories were examined by computing the mean squared displacements (MSD) and the velocity autocorrelation functions (VAF) (Fig. 3b-d). MSD and VAF were calculated using pairwise V_H_-D_H_J_H_ and D_H_J_H_-D_H_J_H_ distance trajectories, which eliminated the effect of nuclear motion. Time-averaged and time-and-ensemble-averaged MSD exhibited a subdiffusive scaling exponent (α < 1) both for intrachromosomal V_H_-D_H_J_H_ and interchromosomal D_H_J_H_-D_H_J_H_ motion (Fig. 3b and 3c, Supplementary Fig. 3 and Supplementary Table 1). Specifically, V_H_-D_H_J_H_ motion was found to be extremely subdiffusive, characterized by a scaling exponentα that decreased from 0.35 at short time scales down to 0.2 at long time scales (Fig. 3c and Supplementary Table 1). VAFs exhibited negative correlations indicative of a push-back from the environment (Fig. 3d). For the intrachromosomal motion, the value of VAF at the dip approached the theoretical limit (−0.5) of an extreme confinement (Fig. 3d). The collapse of the velocity autocorrelation curves upon rescaling of the time axis indicated that the motion was self-similar, i.e. exhibited similar patterns on different spatial and temporal scales (Fig. 3d, right panel). Taken together, these data indicate that V_H_-D_H_J_H_ motion is extremely subdiffusive as the result of a viscoelastic and spatially confined environment.

### Global confinement from chromatin loops eliminates bias in variable region usage

To explore the mechanistic origin of the viscoelasticity and confinement that govern V_H_-D_H_J_H_ motion, we modeled the chromatin fiber as a bead-spring polymer chain using Molecular Dynamics (MD) simulations (Fig. 4)^24^. Given the large number of molecular components and an extensive parameter space that could in principle be incorporated in the simulations, we constrained the model by utilizing multiple independent sets of experimental data. As the first step, we sought to identify the minimal model that could reproduce the structural properties of the Igh locus, namely the plateau in the V_H_-D_H_J_H_ spatial distances as a function of the genomic distance observed in 3D FISH^25,26^. As the second step, which is described in the next section, the resulting minimal model was refined such that it could, in addition, reproduce the dynamic properties of the locus, namely the de-mixing of the V_H_-D_H_J_H_ distances and the strongly subdiffusive scaling exponent.

**Fig. 4.**
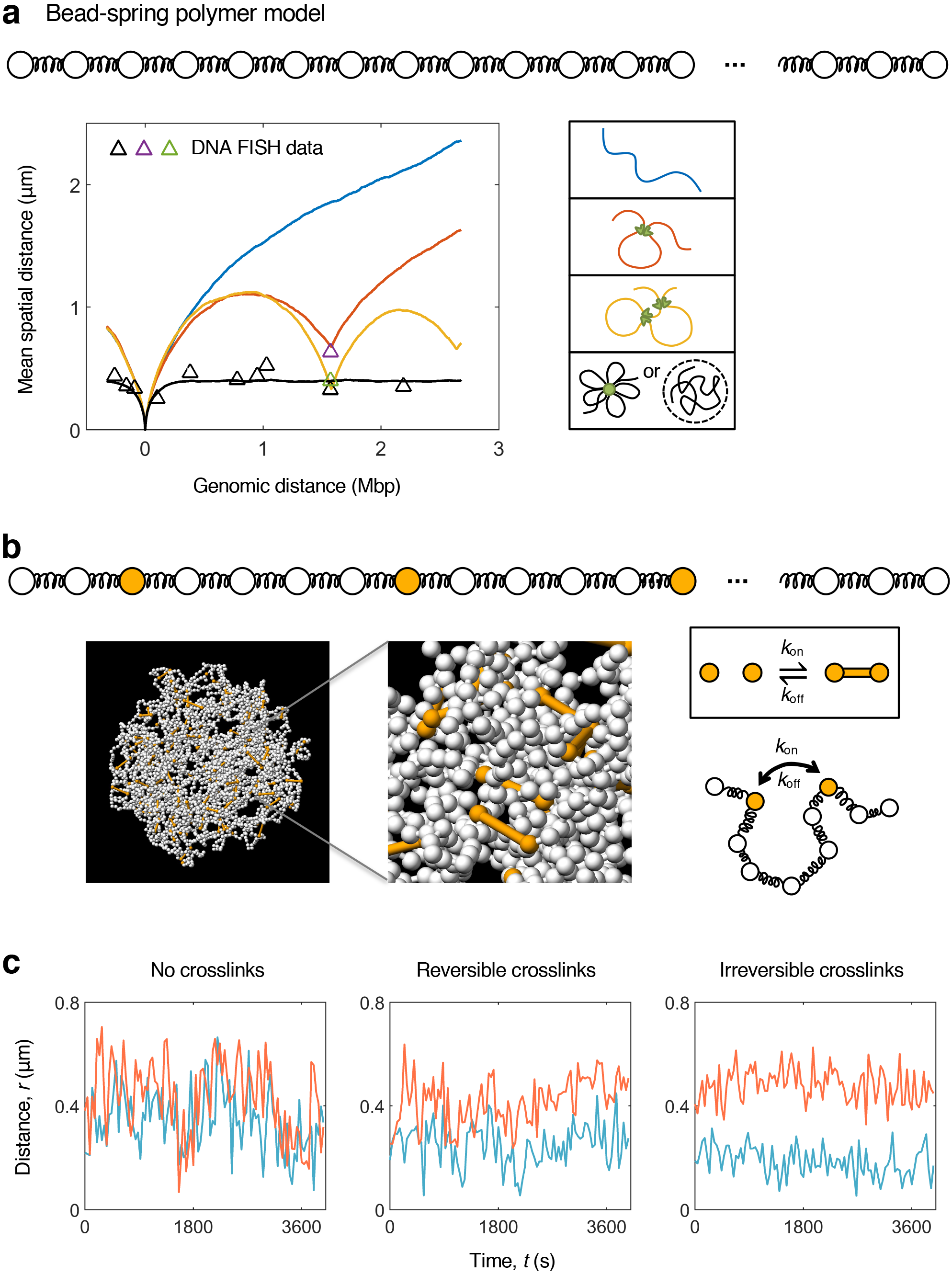
Molecular dynamics simulations capture structural and dynamic properties of the Igh locus. **a,** The mean spatial separation as a function of the genomic distance between the D_H_J_H_ element and the V_H_-repertoire from FISH measurements (triangles) and from MD simulations of a hierarchy of polymer configurations (lines). A multiple-loop chromatin configuration, or, equivalently, a global spatial confinement, reproduces the experimentally observed plateau. **b,** A snapshot from MD simulations of the bead-spring model of the Igh locus with cross-linkable sites. **c,** The insertion of crosslinks across the Igh locus in the simulations reproduces the experimentally observed de-mixing of the trajectories.

To capture the structural properties of the Igh locus, we simulated a hierarchy of worm-like chain polymer models reflecting chromatin configurations of increasing complexity: an unstructured chromatin chain, a single loop configuration, a two-loop configuration, and a chain enclosed in a confinement that mimicked a multiple-loop configuration averaged over many realizations of different ways of looping (Fig. 4a). As expected, the unstructured chain model failed to generate the plateau in the spatial versus genomic distances, instead producing a monotonically increasing dependence (Fig. 4a). A spatial confinement enclosing the chain was necessary to achieve the saturation of the spatial distance at 0.4 μm over more than 2 Mb of genomic distances as observed in 3D-FISH experiments^25,26^. Similarly, the average over a large number of explicit multiple-loop configurations in the absence of confinement also reproduced the plateau (Supplementary Fig. 4). These results indicate that chromatin loops act as a primary source of large-scale confinement, which enables V_H_ regions that are spread over vast genomic distances to reach the D_H_J_H_ region with similar frequencies, independent of their genomic proximity to the D_H_J_H_ region.

### Immunoglobulin heavy chain locus in a gel-like state

While the model of the Igh locus compartmentalized as loop domains correctly reproduced the structural properties (Fig. 4a), it yielded intersecting dynamic trajectories (Fig. 4c, left panel) that were inconsistent with the de-mixed trajectories observed in live-cell imaging (Fig, 3a and Supplementary Fig. 2). This discrepancy suggested yet another layer of constraint, which confined the motion locally. A plausible mechanism of such a local confinement is the cross-bridging of the chromatin fiber via cross-links, analogous to the mechanism that was recently proposed for the formation of super-enhancer whereby cross-links induce phase separated condensates^4^. To examine a potential role of cross-links as the source of local confinement, we introduced in the simulations 5% cross-linkable sites across the Igh locus which could undergo pairwise binding and unbinding with tunable kinetics (Fig. 4b; Supplementary Information). Notably, in the presence of crosslinks, the de-mixing of the trajectories was recovered and became progressively pronounced upon increasing cross-link residence time (Fig. 4c).

In polymer physics, a solution of free (un-crosslinked) chains and a network of cross-linked chains are known to represent two distinct physical phases: a sol phase and a gel phase, respectively^27^. The modeling approach described above and guided by the experimental observations both at the population-and single trajectory-levels suggested that the phase state of the Igh locus corresponds to a gel, or a dynamic three-dimensional network of chromatin chains and associated factors, held together by cross-links^28^. As a consequence of gelation, those V_H_ and D_H_J_H_ segments that end up in close spatial proximity by virtue of their chromatin configuration will remain in close proximity and likely encounter each other within a short time (seconds to tens of minutes). Conversely, those V_H_ and D_H_J_H_ segments that end up spatially remote as a result of their chromatin configuration will remain separated for a long time (more than an hour). Thus, gelation, by constraining the motion locally via cross-links, either facilitates or hinders genomic encounters, depending on the initial spatial separation imposed by a particular fold of the chromatin backbone.

Gel is a solid^27^, and thus cross-linking imparts solid-like properties to the locus, which otherwise (in the absence of crosslinks) would possess the liquid-like properties of a sol. Given that the elasticity of the solid phase and the fluidity of the liquid phase can both be advantageous from a biological prospective, we asked: to what extent are the properties of Igh locus solid-like as opposed to liquid-like? Or, equivalently, how deeply in the gel phase is the locus positioned on a phase diagram? The tuning of the bond lifetime of the simulated cross-links from irreversible bonds (strong gel) to short-lived bonds (weak gel) and further to the absence of cross-linking (sol) resulted in a systematic change in the slope (α) of the MSD (Fig. 5a and Supplementary Fig. 5). The bond lifetime on the order of 10 seconds was found to yield the best agreement between simulated and experimentally measured MSD, indicating that the state of the locus corresponds to a weak gel poised close to the boundary of sol and gel phases (Fig. 5a). Taken together, these results indicate that a large-scale confinement imposed by chromatin loops and a local confinement imposed by the cooperative action of cross-links together constitute the mechanistic origin of the strongly subdiffusive V_H_-D_H_J_H_ motion, and that the Igh locus is found in the state of a weak gel.

**Fig. 5.**
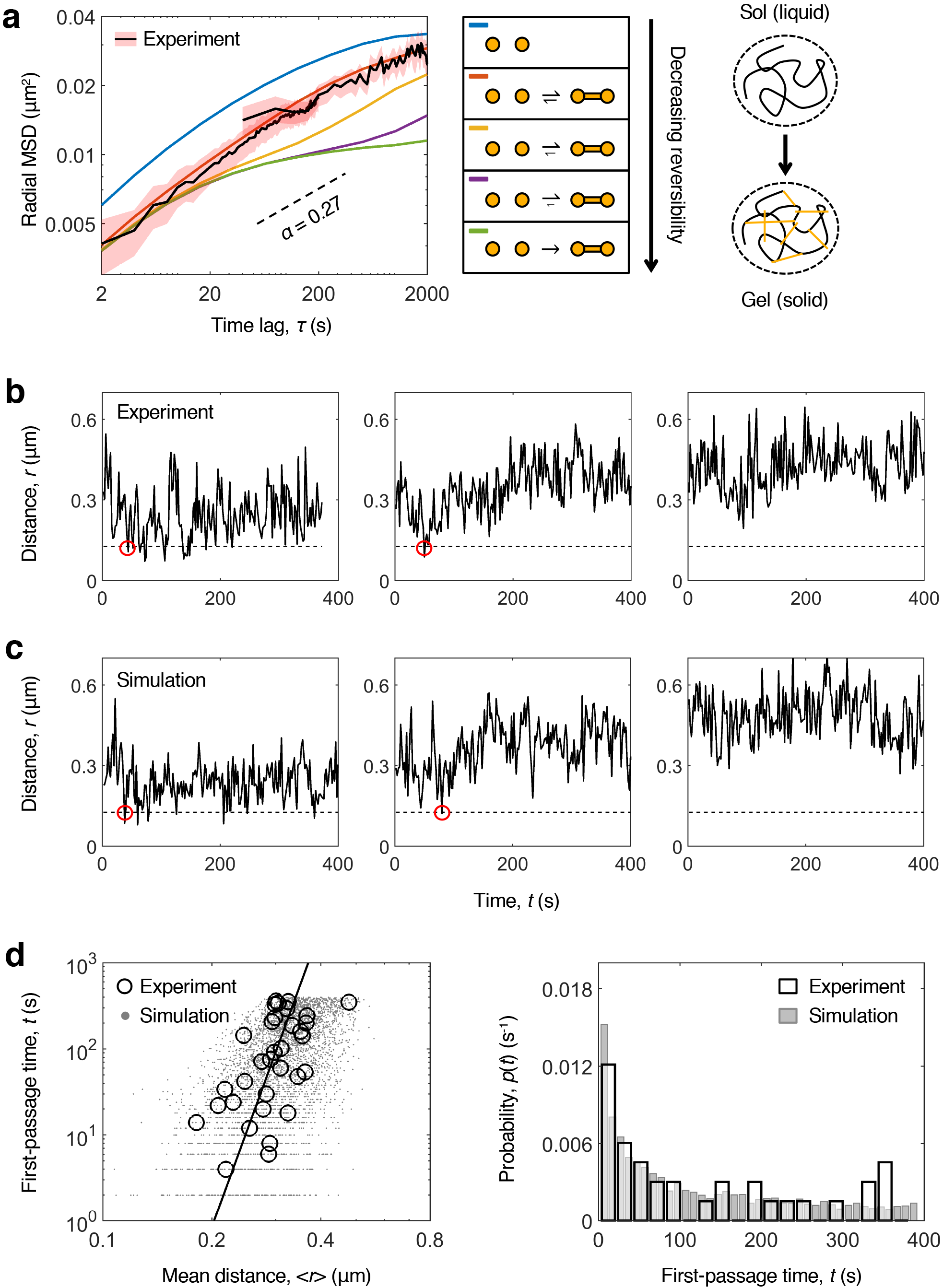
Modeling the Igh locus as a weak gel reproduces V_H_-D_H_J_H_ first-encounter times. **a,** Tuning the lifetime of crosslinks results in a systematic change in the slope (α) of the MSD. Short-lived bonds (lifetime ~ 1 min) yield the best agreement between MSD from experiment and simulations. **b**, Representative trajectories of the spatial distance within individual V_H_-D_H_J_H_ pairs in STI-571 treated cells. Red circles indicate potential first encounters. The corresponding encounter times are within an hour for V_H_ and D_H_J_H_ segments that were in close spatial proximity at the start of imaging. **c**, Representative trajectories of V_H_-D_H_J_H_ spatial distances from the MD simulations of the Igh locus modeled as a weak gel. **d**, First encounter times as a function of mean spatial separation from experimental and simulated V_H_-D_H_J_H_ trajectories (left). The distributions of the first-encounter times from experiment and simulation (right).

### V_H_-D_H_J_H_ encounter times in a gel state

The two-color imaging strategy allowed us to measure the spatial distance within the individual V_H_-D_H_J_H_ pairs as a function of time and thus enabled a direct estimation of the first-passage times (FPTs), or the waiting times of the first encounters (Fig. 5b and Supplementary Fig. 6). We set a threshold value for the distance at which the segments can be considered interacting by combining the value of 30 nm for the true interaction distance and the contribution from localization error (Supplementary Information). A dip in the distance below the threshold indicates a potential encounter, and the corresponding time provides an estimation for the FPT. In order to acquire a rich statistics of the encounter events, we generated a pool of STI-571 treated pro-B cells, which yielded a significantly higher fraction of potential encounters. In the sub-population of the cells in which V_H_ and D_H_J_H_ segments were spatially remote at the start of imaging (initial separation > 0.55 μm), no potential encounters were detected throughout the imaging period. In contrast, the sub-population of cells with V_H_ and D_H_H_h_ segments spatially co-localized at the start of imaging (< 0.55 μm), the potential encounters occurred already within minutes in more than 40% of these cells (Fig. 5b and Supplementary Fig. 6). To examine whether the weak-gel model of the Igh locus reproduces the experimentally observed first-encounter times, we generated the trajectories of V_H_-D_H_J_H_ distances from MD simulations of a globally confined, weakly cross-linked chain. The range of the first-encounter times extracted from the simulated trajectories (Fig. 5c) was in quantitative agreement with the range observed experimentally (Fig. 5b). To investigate the dependence of the FPTs on the spatial distance, we examined both the measured and simulated FPTs as a function of mean spatial distance, the parameter that sets the length scale of this first-passage-time problem. The experimental and simulated data showed good quantitative agreement (Fig. 5d). Furthermore, on a log-log plot, both types of data resulted in the best fit by a line with a slope of 2/*α* (Fig. 5d and Supplementary Fig. 7), in agreement with the predicted scaling MFPT ~ *R*^2/α^ ^19^. Taken together, the first-encounter times for V_H_-D_H_J_H_ interactions from live cell imaging, as well as the scaling of these times with the V_H_-D_H_J_H_ spatial distance, were found to be in quantitative agreement with the encounter times generated by a model of the Igh locus as a cross-linked network of chromatin and associated factors that is poised near the sol/gel phase boundary.

## Discussion

It is now established that the Igh locus is organized into loop domains (1,2). However, our understanding of the mechanisms that govern the dynamics of V(D)J rearrangement remains rudimentary at best. The approach developed here enabled us to simultaneously track the motion of V_H_ and D_H_J_H_ elements and to visualize V_H_-D_H_J_H_ encounters in live B cells. The resulting findings highlight the link between the spatial and temporal aspects of Igh locus organization. Namely, the interplay between the confinement imposed by loop domains and crosslinks on the one hand and the inherently subdiffusive motion of the chromatin fiber on the other hand can either facilitate or hinder genomic interactions within the Igh locus. Indeed, in mediating genomic encounters, subdiffusive V_H_-D_H_J_H_ motion has its advantages and disadvantages compared to normal diffusion: a sub-diffuser searches more efficiently at short distances due to its tendency to explore the space compactly, but it is less efficient searching over long distances due to the slower growth of its MSD with time. Spatial confinement enhances each of these two tendencies: V_H_ and D_H_J_H_ segments that are brought into spatial proximity through looping and whose proximity is further reinforced by cross-links become spatially co-localized; this amplifies the efficiency of their subdiffusive search over short distances, thereby facilitating their encounters. Conversely, V_H_ and D_H_J_H_ gene segments that end up in different loops, which are further stabilized by crosslinks, become isolated from each other; this amplifies the inefficiency of their subdiffusive search over long distances, further hindering their encounters.

Our study further revealed that the structural and dynamic properties of the Igh locus are consistent with those of a weak gel, or a network of chromatin chains and associated factors that are bridged via short-lived cross-links. We concluded that, in the phase space, the Igh locus is positioned within the gel phase (and thus it is a solid) but close to the sol/gel (or liquid/solid) phase boundary^29^, akin to a molded pudding with barely enough gelatin to hold its form. Given that the lifetime of cross-links in the cell is regulated by epigenetic marks, non-coding transcription and chromatin remodelers, and thus represents a dynamical quantity, our findings suggest that the gelation of chromatin is a biological process that is finely tuned with respect to a dynamical variable – the strength of the cross-links. Proximity to the liquid/solid phase boundary allows the loop domains within the Igh locus to rapidly switch their chromatin state by tuning the deposition of epigenetic marks and recruiting chromatin remodelers. Tuning the chromatin properties further into the solid phase (gel) promotes the formation of phase-separated gel droplets, which in turn facilitate the encounters between spatially close D_H_ and J_H_ (or V_H_ and D_H_J_H_) segments and hinder the encounters between spatially remote segments. Conversely, tuning the chromatin state closer to the liquid phase (sol) allows the existing gel droplets to dissolve and new gel droplets to form readily, thus enabling a rapid and ordered re-assembly of the Igh locus.

On the basis of these findings, we propose that the epigenetically-controlled formation of gel droplets comprising chromatin loop domains introduces two modes in V_H_-D_H_J_H_ and D_H_-J_H_ encounters – facilitation and hindrance, thereby preventing premature rearrangements and enforcing VDJ recombination to proceed in a stepwise fashion. At the same time, the proximity of the Igh locus to the sol/gel phase boundary and the resulting propensity to switch between liquid and solid states enables a rapid re-assembly of the locus such that the VDJ recombination can readily proceed from one step to the next. Weak gelation therefore offers a tradeoff between stability and responsiveness in the generation of antigen receptor diversity.

Finally, what is the molecular nature of the cross-links that result in gelation of the Igh locus? It has long been established that the chromatin landscape across the V_H_ and D_H_J_H_ regions is developmentally regulated^13^. Immediately prior to D_H_-J_H_ rearrangement, the D_H_-J_H_ loop domain is closely associated with non-coding transcription, and histones across the D_H_-J_H_ loop domain are hyperacetylated^20,30^. Likewise, the onset of V_H_-D_H_J_H_ rearrangement is closely associated with non-coding transcription, epigenetic modifications and the recruitment of chromatin remodelers across the V_H_ region^17,31,32,33^. It has been widely assumed that these developmentally regulated alterations of chromatin across the Igh locus simply function to promote accessibility^32^. Here we propose that epigenetic modifications of the chromatin fiber across the Igh locus generate, in a developmentally regulated fashion, a dispersed population of weak, reversible cross-links, which, in turn, induce phase-separated gel droplets, or segregated regions of cross-bridged chromatin fiber (Fig. 6)^4,34^. Such cross-links would first induce a gel droplet comprising the D_H_-J_H_ loop domain, thereby facilitating D_H_-J_H_ encounters within the loop domain and simultaneously functioning as an insulator to prevent the D_H_ and J_H_ segments from prematurely interacting with V_H_ segments located outside the D_H_-J_H_ loop domain. Upon the formation of the D_H_J_H_ segment, epigenetic marks would be erased and non-coding transcription halted to allow the cross-links to dissociate and the gel droplet to dissolve. This step would lead to the disassembly of the D_H_J_H_ loop domain, formation of a *de novo* loop domain containing V_H_ and D_H_J_H_ elements, induction of epigenetic modifications and non-coding transcription across the V_H_ region, the formation of a phase-separated gel droplet comprising the V_H_-D_H_J_H_ loop domain, and culminating in the V_H_-D_H_J_H_ rearrangement. In conclusion, we propose that gelation within chromatin loop domains, induced by epigenetic modifications, chromatin remodelers and non-coding transcription, introduces the facilitation and hindrance modes in V_H_-D_H_J_H_ and D_H_-J_H_ encounters thereby orchestrating a stepwise, ordered process for antigen receptor rearrangement. At the same time, the proximity of the Igh locus to the sol/gel boundary in the phase space, and the resulting propensity to switch between solid and liquid states, offers a tradeoff between stability and responsiveness in the generation of antigen receptor diversity.

**Fig. 6.**
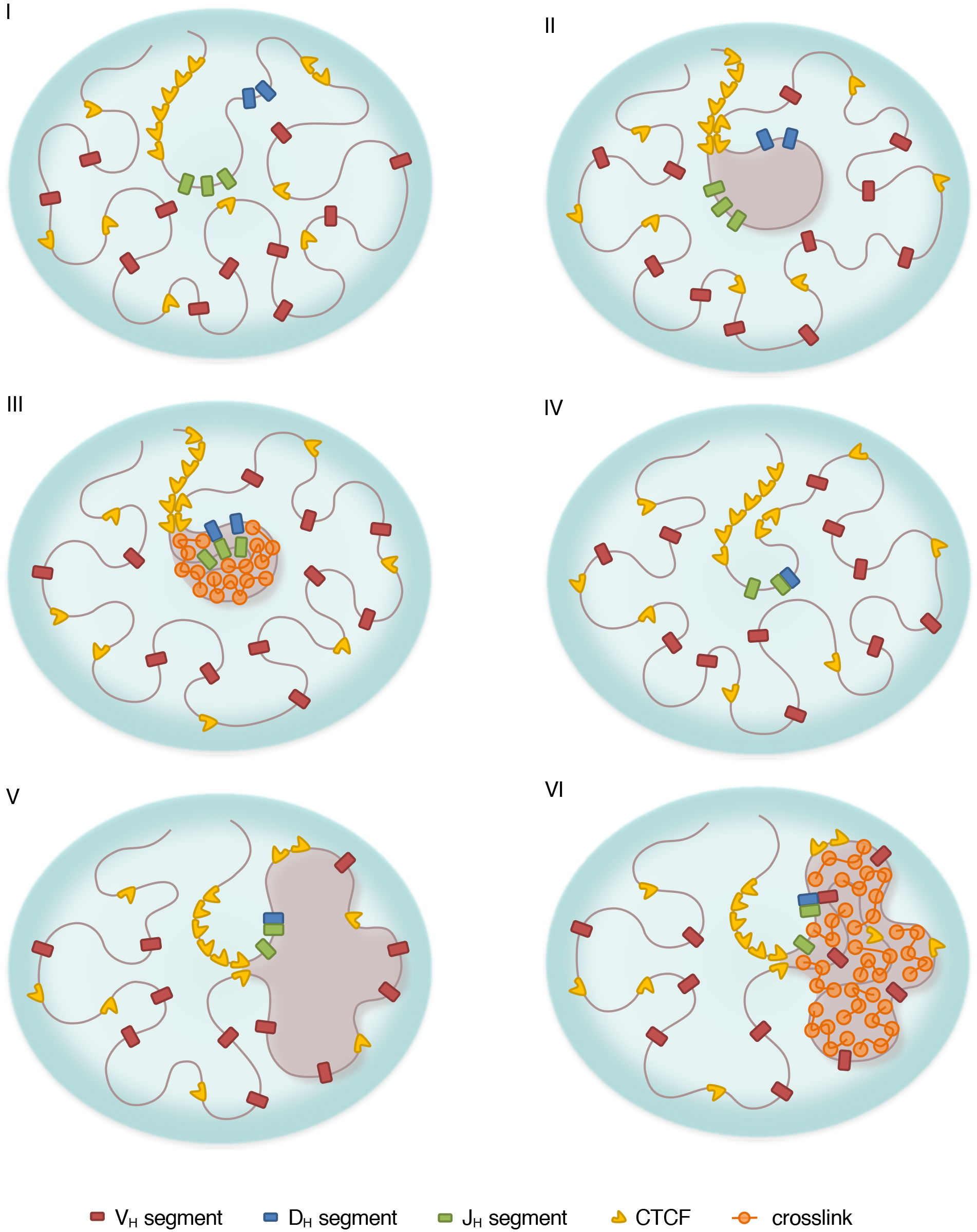
V(D)J Recombination orchestrated by gelation and proximity to the sol-gel phase transition. (I) Igh locus harboring D_H_, J_H_ and V_H_ regions in a germ-line configuration, with no loop domains or cross-links. (II) A chromatin loop domain containing D_H_ and J_H_ regions is formed, anchored by the CTCF sites flanking the D_H_ elements and the CTCF sites in the super-anchor. (III) Weak crosslinks are established across the loop domain containing D_H_ and J_H_ regions. The resulting gel-droplet, which is phase-separated from the rest of the locus, facilitates encounters between D_H_ and J_H_ elements and hinders encounters involving V_H_ elements. (IV) The CTCF sites are inactivated and crosslinks are dissolved; the mobility of the newly formed D_H_J_H_ segment is increased. (V) Chromatin loops containing D_H_J_H_ and V_H_ regions are formed, anchored by the CTCF sites across the V_H_-cluster and the CTCF sites in the super-anchor. (VI) Cross-links across a loop are established, resulting in a new phase-separated gel-droplet that facilitates encounters between spatially close V_H_ and D_H_J_H_ elements.

## Methods

### Construction of tandem arrays of MUT-TET repressor binding sites

WT-TET repressor binding sites were composed of TCTCTATCAGTGATAGGGA DNA sequences. MUT-TET repressor binding sites contained the TCTCCGTCAGTGACGGGGA DNA sequence. DNA fragments containing four randomly repeated TET-repressor binding sites, each flanked by 10 bp random DNA sequence, were generated from synthetic oligonucleotides and inserted into a plasmid backbone using Nhe1, Xba1 and Pst1 restriction sites. Tandem arrays containing 320 copies of the MUT-TET binding sites were generated by sequential cloning and expansion of the number of repeats as described previously (Lau et al., 2003).

### Constructs and Retrovirus Production

To generate Sc-Mut-TetR(B)-SNAP, mutant TetR was generated by site directed mutagenesis of TetR-EGFP to replacing glutamic acid residues at position 37 to alanine and proline at position 39 to lysine (Lucas et al. 2014). The above-generated vector was digested by MfeI and SalI and ligated with a MfeI-and SalI-digested gBlock segment containing SG_4_ linker, Mutant-TetR, and SNAP tag to obtain Sc-Mut-TetR(B)-SNAP. Vectors were purified by CsCl gradient equilibrium centrifugation and transfected into 293T cells along with retroviral packaging vectors by calcium phosphate transfection. Medium was replaced the following morning, and viral supernatant was harvested 1 day later and stored at −80 C until used.

### Cell Culture and Transduction

B lineage cells were grown in OptiMEM media with 10% FCS and 1% Pen-Strep-Glutamine (Life Technologies) in the presence of IL7 and SCF. Supernatant from IL7-expressing cell lines was added at a concentration previously determined to promote B cell growth. Cells were incubated at 37°C under 5% CO2. TetR-EGFP and Sc-Mut-TetR-SNAP were co-infected and after two days cells were labeled with either SNAP-cell 647-SiR or SNAP-cell TMR-Star according to manufacturer’s (NEB) instructions. For the locus contraction experiments, cells were treated with 10 |jm STI-571 for 36 hours before performing 3D DNA FISH or live imaging. 3D DNA FISH was performed as described before (Jhunjhunwala et al.).

### Imaging

Imaging was performed under normal growth conditions using phenol red-free media (Carlton et al., 2010). Cells were plated on Fluorodish poly(D-lysine) coated plates (World Precision Instruments) at least 2 hours before imaging. Cells were imaged using a 100X 1.4NA oil immersion objective on an Applied Precision OMX microscope equipped with an EM-CCD camera. 20-30 Z sections were obtained with a spacing of 0.5 μm. Images were acquired at a rate of one stack every two seconds for a total of 200 total time points or one stack every 40 seconds for a total of 120 time points.

### Imaging Analysis

An Applied Precision OMX microscope was used for live-cell microscopy. Data were deconvolved using Applied Precision SoftWorx software. The center of mass of Igh loci TetO probes was tracked at each time point using the “spots” feature of Bitplane’s Imaris software. The “quality” threshold was manually adjusted for each cell so that the TetO probes were recognized at each time point. The “track duration” filter was used to remove incorrectly recognized spots.

### Modeling the chromatin fiber as a worm-like chain

The 3 Mb Igh locus was modeled as a bead-spring self-avoiding worm-like chain, using an open source Molecular Dynamics simulation platform LAMMPS (Plimpton, 1995). The chromatin fiber was simulated using, as parameter values, 50 bp/nm for the packing density of the fiber and 14 nm for the diameter of the bead, resulting in 700 bp per bead and a total of 4298 beads in the Igh locus. These values are consistent with estimates derived from experiments (Jhunjhunwala et al., 2008; Sanborn et al., 2015; Ou et al., 2017). The known genomic locations of individual V_H_, D_H_, J_H_ and CTCF elements determined the beads on which these elements were positioned in the simulations. The consecutive beads were connected by harmonic springs. The repulsive part of the Lennard-Jones potential was used to account for the physical volume occupied by each bead. The choice of parameters allowed occasional chain self-crossing, mimicking a moderate topoisomerase activity. The persistence of the chain was modeled through an angle potential which gave rise to a persistence length of approximately 22 nm. The dynamics of the chain was simulated using LAMMPS fix commands that perform Langevin dynamics according to the Langevin equation.

### Simulating a spectrum of chromatin configurations

The hierarchy of the simulated polymer models (Fig. 5a) included a simple unstructured chromatin chain, a one-loop configuration of the chain, a two-loop configuration, and a spatially confined chain mimicking the effect of an “averaged” multiple-loop configuration. The looped configurations were engineered by introducing harmonic spring-bonds between the beads that anchored the loops. The simulation of a spatially confined chain was performed by introducing a global spherical confining potential of diameter 0.8 μm to enclose the otherwise unstructured chain. To obtain the spatial distance vs genomic distance relation, a statistical ensemble of 1000 chains was generated for each chromatin configuration.

### Simulating crosslinks with varying degree of reversibility

Weak intra-chain interactions were modeled by introducing 5% equidistant crosslinkable sites into the simulation of the spatially confined chain (Fig. 5b; Supplementary Information). The crosslinkable beads could undergo pairwise binding/unbinding events. The degree of reversibility of the crosslinks was varied by tuning the lifetime of the crosslinks. In order to explore the exclusive effect of crosslinks, a control set of simulations was performed by replacing the global confining potential with periodic boundary conditions.

### First-passage time analysis of experimental and simulated V_H_-D_H_J_H_ trajectories

A potential encounter event was identified as a dip in the distance between the V_H_ and D_H_J_H_ segments below a threshold value. The threshold value *r*_c_ was determined by combining the value of the true interaction distance *r*_int_ and the localization error *r*_err_ through *r*_c_ = (*r*_int_^2^ + *r*_err_^2^)^0.5^. The value of *r*_int_ was set to be 30 nm, and the value of *r*_err_^2^ was extracted from the control experiment on measurement error to be 0.015 μm^2^. The time corresponding to the first dip provides an estimation for the first-passage time. The first-encounter times from experiment and simulation as a function of mean separation distance within the V_H_-D_H_J_H_ pair, as well as the distributions of the first-encounter times, were compared (Fig. 5d). The first-encounter times from experiment were fitted with a power-law relationship FPT = *β*<*r*>^2/α^. The fit yielded *α* = 0.17 and *β* = 10^8^ μm^−2/α^s. The value of α is close to α = 0.13 extracted from the fit of the MSD of STI-571 treated pro-B cells, thus validating the predicted scaling MFPT ~ *R*^2/α^.

## Acknowledgements

We thank Dr. Peter Geiduschek for insightful comments, suggestions and editing. We thank the members of the Murre laboratory and the Dudko group for stimulating discussions. We thank Jennifer Santini for help with imaging and image analysis. We thank Dr. William Bialek for insightful discussions. We thank Dr. Ned Wingreen for valuable insights and advice relating to the simulations. Our studies were supported by grants from the National Institutes of Health (U54DK107977) to O.K.D. and the National Institutes of Health (U54DK24230, AI082850, AI00880 and AI09599) to C.M. Y.Z. was supported by the Princeton Center for Theoretical Science the National Science Foundation (Grant PHY1607612), and the NSF Center for the Physics of Biological Function (PHY1734030).

## Authors contributions

N.K. generated tandem arrays of mutant TET-repressor binding sites, cell lines that harbored MUT-TET operator binding sites, performed the live cell imaging experiments and measured the spatial distances as a function of elapsed time. Y.Z. analyzed the data, designed and performed simulations. J.S.L generated pro-B cell lines carrying tandem arrays of TET-operator binding sites. O.K.D. and C.M. supervised the study and wrote the paper. All authors discussed the results and commented on the manuscript.

## Integrated supplementary information

**Supplementary Figure 1. Extracting genomic motion from observed motion.** The relative motion of the genomic segments was extracted from the observed motion by eliminating the effect of nuclear rotation and the measurement error. To ensure that the extracted motion of the genomic segments was unaffected by the extraction procedure, two independent approaches, both involving no adjusting parameters, were utilized and demonstrated to yield nearly identical results across short (left) and long (right) time scales.

**Supplementary Figure 2. Tracking V_H_-D_H_J_H_ Motion in Live B Cell Progenitors Reveals a Spectrum of Dynamic yet Stable Chromatin Configurations.** Temporal trajectories of V_H_-D_H_J_H_ spatial distances, color-coded according to their mean values, reveal a de-mixing effect, visualized as the “rainbow” pattern, whereby the distances for the individual V_H_-D_H_J_H_ pairs fluctuate around their respective mean values that remained nearly constant for at least an hour.

**Supplementary Figure 3. Mean squared displacement of inter-chromosomal motion is subdiffusive.** Time-averaged radial MSD (colored lines) and the time-and-ensemble-averaged radial MSD (black line) for the inter-chromosomal (D_H_J_H_-D_H_J_H_) motion in pro-B cells. Both types of MSD are subdiffusive (α < 1) and are characterized by the similar values of the anomalous scaling exponent α (the slope) and the anomalous diffusion coefficient *D* (the vertical offset). The short horizontal black line indicates the mean measurement error.

**Supplementary Figure 4. The mean spatial distance between D_H_J_H_ and V_H_ averaged over 1000 explicit eight-loop configurations reproduced the experimentally observed plateau.** Individual eight-loop configurations were engineered by anchoring the CTCF site at the superanchor to 7 randomly selected CTCF sites in the V_H_ region and the most distal CTCF site. The mean spatial distance as a function of the genomic distance was computed from the ideal-chain model for individual configurations (gray lines) and averaged over all random looping configurations (black line).

**Supplementary Figure 5. The exclusive effect of crosslinks on the genomic motion.** The time-averaged radial MSD (gray lines) and time-and-ensemble-averaged radial MSD (colored lines) of the relative motion between the marked V_H_ and D_H_J_H_ elements in the set of simulations with varying degree of reversibility of crosslinks and with the periodic boundary condition.

**Supplementary Figure 6. Representative V_H_-D_H_J_H_ trajectories across long time-scales.** Potential first-encounter events are marked with red circles. The estimated FPTs are on the timescale of seconds to hours for the V_H_ and D_H_J_H_ elements that were found in a close spatial proximity at the start of imaging.

**Supplementary Figure 7. Mean spatial distances and mean first-encounter times from the molecular dynamics simulations for a hierarchy of the chromatin topologies. a**, Chromatin loops bring genomically distant segments to spatial proximity. **b**, Looped configurations of chromatin reduce the mean first-encounter times between the D_H_J_H_ element and V_H_ elements from tens of hours (*left*) down to biologically relevant timescales (seconds to a few hours, *right*). **c**, Mean first-encounter times as a function of the mean V_H_-D_H_J_H_ spatial distance for the hierarchy of chromatin topologies. The best fit results in the slope of 4, confirming the predicted scaling MFPT ~ *R*^2/α^(*α* = 0.5 in the simulations).

## Theory and Modeling

### Extracting genomic motion from observed dynamics

The observed apparent relative motion of the genomic segments is a result of a superposition of the true relative motion of the genomic segments, the rotation of the cell and nucleus, and measurement error. We developed two independent procedures to extract the true genomic motion from the observed apparent dynamics. In the first procedure, we eliminated the rotational motion by taking advantage of the availability of pairwise V_H_-D_H_J_H_ and D_H_J_H_-D_H_J_H_ spatial distances, which are unaffected by rotation, and performing the MSD analysis on the distance trajectories *r*(*t*) as MSD = <*r*(*t*) − *r*(*t* + τ))^2^>. As the changes in distances reflect the changes in the radial direction of the true relative motion in three dimensions (3D), we refer to the MSD calculated this way as “radial MSD”. It can be shown that, under the condition |*r*(*t*) − *r*(*t* + *τ*)| ≪ *r*(*t*), the radial MSD from the observed distance trajectories, radMSD^observed^, is equal to 1/3 of the MSD of the true relative motion in 3D, MSD^genomic^, plus measurement error, MSD^err^. The factor 1/3 reflects the fact that the radial motion represents 1 out of 3 degrees of freedom of the full 3D motion. The MSD of the genomic motion between segments in 3D is then

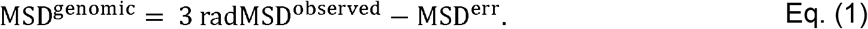

Eq. (1) gives the MSD of the true relative motion for each of the two pairs of genomic segments, the intra-chromosomal V_H_-D_H_J_H_ pair and the inter-chromosomal D_H_J_H_-D_H_J_H_ pair. We assessed the contribution of the measurement error, MSD^err^, by performing a control experiment in which a red and a green markers were inserted in the same genomic region, so that any observed relative motion between these markers would be due to measurement error only. The MSD^err^ was calculated using the standard formula, (***r***(*t*) − ***r***(*t* + *τ*))^2^>, where ***r***(*t*) is the displacement vector between the two markers in the control experiment. We found the value of the MSD^err^ to be approximately 0.02 μm^2^, or an average localization error of 22 nm for the *x*- and *y*- coordinates and 95 nm for the z-coordinate for each marker. The MSD plots in Fig. 4 in the main text were obtained following the above procedure.

In the second procedure, we used the fact that the genomic motion can be obtained by subtracting all other contributions away from the observed motion:

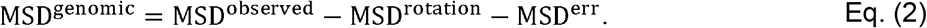

The separation of individual contributions is valid as long as there is no correlation among the different types of motion and measurement error. Eq. (2) applies both to the V_H_-D_H_J_H_ and D_H_J_H_-D_h_J_h_ pairs. The MSD^observed^ can be calculated directly from experimental data, MSD = <(***r***(*t*) − ***r***(*t* + *τ*))^2^>. The MSD^err^ was assessed through the measurement error control experiment described in the first procedure. The rotational contribution MSD^rotation^ for the V_H_-D_H_J_H_ pair was estimated as follows. By substituting Eq. (1) into Eq. (2), we found:

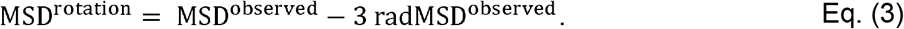

As each of the terms on the right-hand side can be obtained directly from the experimental data, we used Eq. (3) to evaluate the rotational contribution MSD^rotation^ of the D_H_J_H_-D_H_J_H_ pair. We then utilized MSD^rotation^ of the D_H_J_H_-D_H_J_H_ pair to estimate MSD^rotation^ of the V_H_-D_H_J_H_ pair by noting that, on average, the amplitude of the rotational motion of a pair of segments is proportional to the mean distance between the segments in that pair:

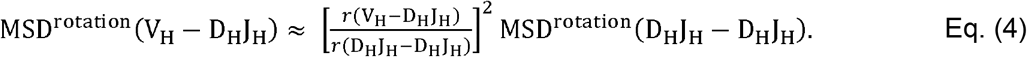

Here, *r*(V_H_-D_H_J_H_) and *r*(D_H_J_H_-D_H_J_H_) are the mean distances between the segments of the corresponding pairs. We then obtained the true relative genomic motion of the V_H_-D_H_J_H_ pair by substituting the MSD^rotation^ of the V_H_-D_H_J_H_ pair found from Eq. (4) into Eq. (2).

Finally, as a self-consistency check, we compared the genomic motions of the V_H_-D_H_J_H_ pair extracted via these two procedures, Eq. (1) and Eq. (2). The two procedures yielded nearly identical results for the entire imaging time (Supplementary Figure 1), indicating that the extracted genomic motion is independent of the procedure used. Note that all the terms in Eq. (1) and Eq. (2) were calculated from the experimental data, and therefore both approaches involve no adjustable parameters.

### Modeling the chromatin fiber as a worm-like chain

The mouse immunoglobulin heavy chain locus is located on chromosome 12, spanning the 3 Mb region between nucleotides 117,349,200 and 114,341,024. We modeled the Igh locus as a bead-spring self-avoiding worm-like chain, using an open source molecular dynamics simulation platform LAMMPS^1^. The parameter values for the simulated chromatin fiber were chosen by combining estimates reported in multiple experiments^2,3,4^. Specifically, we assumed the packing density of chromatin fiber to be 50 bp/nm and the diameter of the bead (*d*) to be 14 nm, leading to 700 bp per bead and a total of 4298 beads in the Igh locus. The known genomic locations of individual V_H_, D_H_, J_H_ and CTCF elements determined the beads on which these elements were positioned in the simulations. Specifically, the fluorescently marked V_H_ and D_H_J_H_ regions were positioned on beads 2712 and 467, respectively. The consecutive beads were connected through the harmonic spring potential *k*(*r* − *d*)^2^ with the spring constant *k* = 25*k*_B_*T*/*d*^2^, where *k*_B_ is the Boltzmann constant and *T* was set to the room temperature, 300 K. To account for the physical volume occupied by each bead, we introduced repulsive interactions between all pairs of beads except neighboring beads. The repulsion was modeled using the Lennard-Jones potential, 4*ɛ*[(*σ*/*r*)^12^ − (*σ*/*r*)^6^], with *ɛ* = 1*k*_B_*T* and *σ* = *d*/1.12. The potential was cut off and shifted to 0 at *r* = *d* so that only the repulsive part of the potential was used in the simulation. The above choice of parameters allows occasional chain self-crossing, mimicking a moderate topoisomerase activity. The persistence of the chain was modeled through the angle potential a·cos(*θ*) with *θ* being the angle formed between three successive beads. A proper choice of the prefactor in the bending energy, a = 2*k*_B_*T*, resulted in a persistence length of approximately 22 nm.

The dynamics of the chain was simulated using the commands fix nve and fix langevin in LAMMPS, which performs Langevin dynamics according to the Langevin equation: *md*^2^***R***_*i*_/*dt*^2^ = − *ξ d****R***_*i*_/*dt* − *dU*/*d****R***_*i*_ + ***f***. In the equation, *m* is the mass of the bead, ***R***_*i*_ is the center position of the bead *i*, *ξ* is the friction coefficient, *U* includes all the interaction potentials described above, and ***f*** is the Gaussian-distributed noise with <*f*_α_(*t*) *f*_β_(*t’*)> = 2*k*_B_*Tξ* δ_αβ_δ(*t* − *t’*) where *α* and *β* denote the *x*-, *y*-, or *z*-component of ***f***. The mass of the bead was estimated by assuming that the density of chromatin fiber is comparable to that of water. The friction coefficient ξ in the damping force term was calculated from Stokes-Einstein relation, *ξ* = 3π*ηd*, where the dynamic viscosity *η* was taken to be 1 Pa s (close to that of honey). Note that even though the reported viscosity of nucleoplasm is much lower, the crowded environment of the nucleus and non-specific adhesive interactions between molecules can result in a much higher effective viscosity for relatively large objects, such as the labeled chromatin segments. The above value of viscosity was chosen to best match the dynamical properties of segment motion observed in our experiments. To speed up the simulations, a larger value of the bead mass was used while keeping *ξ* (or the diffusion coefficient) constant, which led to an increased velocity relaxation time and thus allowed a larger integration time step in the simulation. This method did not affect the accuracy of the simulations as the timescale of interest was much longer than the increased velocity relaxation time, i.e. the simulations were performed in the overdamped regime.

### Simulating a spectrum of chromatin configurations

The hierarchy of the simulated polymer models included a simple unstructured chromatin chain, a one-loop configuration of the chain, a two-loop configuration, and a spatially confined chain mimicking an “averaged” multiple-loop configuration. The one-loop and two-loop configurations were engineered by introducing harmonic-spring bonds between the beads that anchored the loops, using the same bond potential as the harmonic potential used for the springs connecting consecutive beads. Specifically, in the one-loop configuration, the bond was introduced between a CTCF site in the superanchor region (bead 171) and a CTCF site upstream of the marked V region (bead 2723), thus enclosing the marked V_H_ and D_H_J_H_ regions in the same loop. In the two-loop configuration, one bond was introduced between a CTCF site in the superanchor region (bead 171) and the most distal CTCF site (bead 4241), and another bond between the CTCF site adjacent to the marked D_H_J_H_ region (bead 550) and the CTCF site upstream of the marked V_H_ region (bead 2723), thus separating the marked V_H_ and D_H_J_H_ regions in different loops. The simulation of a spatially confined chain was performed by introducing a global spherical confining potential *k*(*d* − *r*)^2^ to enclose the otherwise unstructured chain. Note that *r* in the potential is the distance from the bead to the boundary of the sphere. The potential was only switched on when the bead was within a distance *d* from the boundary, *r* < *d*. The spring constant *k* was set to be the same as that of the springs connecting beads. The diameter of the confining sphere was set to be 0.8 μm.

The spatial distance vs genomic distance relation was obtained from a statistical ensemble of 1000 chains for each chromatin configuration. The spatial distances between the D_H_J_H_ bead and the rest of beads on the chain were calculated for each chain, averaged over the ensemble, and plotted as a function of genomic distance (Fig. 5a).

### First-encounter times for the D_H_J_H_ segment and the V_H_-repertoire in looped configurations

The effect of chromatin topology on the first-encounter times was examined quantitatively by performing Molecular Dynamics simulations of the hierarchy of the polymer configurations described above. Here, the packing density of chromatin fiber was set to be 130 bp/nm and the diameter of the bead to be 30 nm, leading to 772 beads in the Igh locus and a persistence length of approximately 50 nm. The marked V_H_ and D_H_J_H_ regions were positioned on beads 487 and 84. In the one-loop configuration, the bond was introduced between CTCF sites on beads 31 and 489. In the two-loop configuration, one bond was introduced between CTCF sites on beads 31 and 762, and another bond between CTCF sites on beads 99 and 489. The interaction distance, which signified a genomic encounter, was set to be 60 nm.

To find the first-encounter time, the distance between V_H_ and D_H_J_H_ beads was checked once every time interval equal to the velocity relaxation time of the beads. Once the D_H_J_H_ bead was within the interaction distance from a V_H_ bead, the corresponding V_H_-D_H_J_H_ encounter time for that V_H_ segment was recorded. To speed up the simulations, a larger value of the bead mass was used. The larger value of the bead mass and the chosen frequency of checking for a potential encounter only had minor effects on the first-encounter times: a control simulation showed that, upon decreasing the bead mass (and hence increasing the checking frequency) by a factor of 100, the first-encounter times were only reduced by a factor of 2. From the simulation, we found that the D_H_J_H_ segment can readily encounter V_H_ segments located on different loops within biologically relevant timescales (Supplementary Figure 7).

### Simulating crosslinks with varying degree of reversibility

The live-imaging data on the Igh locus dynamics indicated that the locus adopts a spectrum of stable configurations. To explore the possibility that weak intra-chain interactions could increase the spatial rigidity of the locus and hence stabilize the locus structure, we introduced 5% equally spaced crosslinkable sites (on beads 10, 30, 50, …) into the simulation of the spatially confined chain (Fig. 5b). The crosslinkable beads could undergo pairwise binding/unbinding events. The bound beads were subjected to a harmonic interaction potential *k*_*b*_(*r* − *d*_*b*_)^2^ with the mean bond length *d*_*b*_ = 50 nm and a soft spring constant, *k*_*b*_ = 0.1*k*. We performed a set of simulations with decreasing degree of reversibility of the crosslinks. The simulation without crosslinks (blue in Fig. 6a) is the same as the original simulation of the spatially confined chain. The simulations with reversible crosslinks were performed using the commands fix bond/create and fix bond/break in LAMMPS. Bonds were checked for creation once every 0.01 s, and were created between unbound crossl-inkable bead pairs when their separations were smaller than 60 nm. The created bonds were checked for breaking once every 1 s, 10 s, or 100 s (orange, yellow, purple in Fig. 6a), and were broken if the separations between the bound beads were larger than 60 nm. We estimated the bond lifetimes in these simulations to be about 10 s, 100 s, and 1000 s, respectively. The simulation with irreversible crosslinks (green in Fig. 6a) was performed by only using the command fix bond/create. To keep the bond information updated, we replaced the original fix_bond_create.cpp file in LAMMPS MC package, which only counts the bonds once at the beginning of the simulation run, with a modified version^7^. Another set of simulations, which aimed to explore the exclusive effect of crosslinks (Supplementary Figure 5), was performed by replacing the global confining potential with periodic boundary conditions using a box size 0.6 μm. Each run in both sets of simulations with different degree of reversibility of crosslinks was repeated 10 times using a different random seed. The *x*-, *y*-, and *z*-coordinates of the crosslinkable beads in the chain were recorded once every 2 s for 4000 s to mimic the experimental protocol. To find the mean squared displacement of the relative motion between the marked V_H_ and D_H_J_H_ beads (2712 and 467), the spatial distance between the two beads was calculated at each time point. The resulting distance trajectories were used in the MSD analysis (Fig. 6a). To enrich the statistics, we included in the MSD analysis the trajectories of the distances between beads with the same genomic separation as that of V_H_ and D_H_J_H_ beads.

### First-encounter-time analysis of experimental and simulated V_H_-D_H_J_H_ trajectories

The two-color imaging strategy enabled a direct estimation of the V_H_-D_H_J_H_ first-encounter times. A potential encounter event was identified as a dip in the distance between the V_H_ and D_H_J_H_ segments below a threshold value. The threshold value *r*_c_ was determined by combining the value of the true interaction distance *r*_int_ and the localization error *r*_err_ through *r*_c_ = (*r*_int_^2^ + *r*_err_^2^)^0.5^. The value of *r*_int_ was set to be 30 nm, and the value of *r*_err_^2^ was extracted from the control experiment by averaging the squared distance between the two markers and found to be 0.015 μm^2^. Note that this value for *r*_err_^2^ is slightly larger than the value expected theoretically, *r*_err_^2^ = 0.5 MSD^err^, likely due to the effect of chromatic aberration in the experiments. As a dip below the threshold is a potential encounter event, the time corresponding to the first dip provides a lower bound for the V_H_ and D_H_J_H_ first-encounter time. On the other hand, as distance trajectories were recorded at a finite time interval, missed encounter events due to finite recording frequency may lead to an overestimation of the observed first-encounter times.

To investigate the relationship between the first-encounter times and the spatial distance, we generated a pool of contracted cells, in which the smaller spatial distances yielded richer statistics of the potential encounter events. Experimental V_H_-D_H_J_H_ trajectories were recorded once every 2 s for 400 s. To best mimic the experimental protocol, simulated V_H_-D_H_J_H_ trajectories were recorded with the same frequency and duration. Measurement errors were randomly drawn from the trajectories obtained in the control experiment and added to the simulated V_H_-D_H_J_H_ trajectories. To enrich the statistics, the trajectories of the distances between beads with the same genomic separation as that of V_H_ and D_H_J_H_ beads were also included in the first-encounter time analysis.

The first-encounter times from experiment and simulation as a function of mean separation distance within the V_H_-D_H_J_H_ pair, as well as the distributions of the first-encounter times, were compared (Fig. 6d). The first-encounter times from experiment were fitted with a power law relationship FPT = *β*<*r*>^2/α^. As the finite length of trajectories (400 s) sets an upper limit for the observed FPTs, the parameters *α* and *β* were found by minimizing the sum of the squares of the distance between individual data points and the fitted line along the x-axis. The fit yielded *α* = 0.17 and *β* = 10^8^ μm^−2/α^s. The value of α is in agreement with the subdiffusive exponent *α* = 0.13 extracted from the fit of the MSD of contracted cells, confirming the predicted scaling MFPT ~ *R*^2/α^.

